# Reports of *Lipoptena fortisetosa* on dogs and in the environment, and evidence of its widespread establishment in Hungary

**DOI:** 10.1101/2025.11.03.686229

**Authors:** Adrienn Gréta Tóth, Attila Bende, Sándor Hornok, Zsombor Wagenhoffer, Balázs Szulyovszky, Viktória Galla, Petra Vöröskői, Gergő Keve

## Abstract

*Lipoptena fortisetosa* is a louse fly of East Asian origin that is considered a potential vector of several pathogenic bacteria and is most commonly associated with deer (Cervidae). The species has been detected in multiple countries in Europe; however, its exact distribution range is unknown. A single individual was detected on a dog at a veterinary clinic in Budapest, Hungary, a country where the presence of *L. fortisetosa* has not yet been confirmed. After acquiring information regarding the recent whereabouts of the dog, targeted louse fly collection with insect nets was performed in a forest in Central Transdanubia. Of the 30 flying, unfed specimens of *Lipoptena* spp., 23 were morphologically identified as *L. fortisetosa*. Following this, louse fly collections have been conducted across Hungary during the fall of 2025. According to these results, *L. fortisetosa* can be considered widespread in the country, and that this parasite can be effectively transported by dogs. The findings draw attention to the potential medical and veterinary significance of the emergence of novel vectors that may have been introduced through animal transport.

## Introduction

*Lipoptena fortisetosa* is a louse fly of East-Asian origin, most commonly associated with deer, (Cervidae), that is likely to have been introduced to Europe with the sika deer (*Cervus nippon*) (Kurina et al., 2019). Based on the available literature data, it is safe to assume that this insect is currently spreading (Andreani et al., 2020; Gałęcki et al., 2021; Yatsuk et al., 2024). Results from other European countries, such as Austria (Jentzsch, 2023), Italy (Andreani et al., 2020, Andreani et al., 2017; Kurina et al., 2019), Czech Republic (Zeman, 2013), Poland (Gałęcki et al., 2021), and Estonia (Kurina et al., 2019) suggest the widespread European distribution of the insect. Until October 2025, *Lipoptena cervi* was the only species of the genus *Lipoptena* reported from Hungary (Egri and Rigó, 2014; Papp, 2001). During the review of this manuscript, another author reported four additional specimens from Eastern-Hungary (coordinates: 47.09, 20.77), in a local journal (Szöllősi-Tóth, 2025). However, these findings were not confirmed by molecular analysis, which would have been beneficial, especially in light of the emergence of recently described species such as *Lipoptena andaluciensis* (González et al., 2024).

While the original host species of the fly, the sika deer (Yatsuk et al., 2024), is not prevalent in Hungary in the wild, one of its greatest populations today (about 100 individuals) is kept at Fehérvárcsurgó game park. At the turn of the 19th and 20th centuries, several exotic game species, including the sika deer (*Cervus nippon*), were introduced to Hungary. The first individuals arrived at the Fehérvárcsurgó estate in 1910. Later reintroductions from the Budapest Zoo and the Soviet Union re-established the population. However, these latter specimens were Dybowski’s sika deer (*C. nippon hortulorum*) (Faragó, 2007; Szabó, 2010).

The aim of this study was to determine whether the presence of a single specimen found in the city centre of Budapest was accidental or originated from an established population within the country.

## Materials and methods

On 2025.08.26, dog (breed: Hungarian vizsla) visited a veterinary clinic in the city central of Budapest, for the purpose of the removal of a previously diagnosed (on 2025.08.12), solitary, well-circumscribed benign fibroadnexal dysplasia (1.2 × 1 cm) developed in an area of chronic inflammation on the right forelimb, as well as for tartar removal. During the routine examination before the surgery, the veterinarians noticed a single, engorged, wingless louse fly (Diptera: Hippoboscidae) on its fur. Other than the presence of a single parasite and the previously diagnosed benign tumor, no relevant lesions or abnormalities were observed on the dog’s integument, and the animal showed no symptoms suggestive of vector-borne pathogen infection. The parasite was sent to the laboratory of the Department of Parasitology and Zoology, University of Veterinary Medicine Budapest for further examination.

The owner of the dog was surveyed for further information:” Has *the dog travelled abroad during the summer? Has the dog visited any forests recently? If yes, where?”*

Based on the owners’ answers, the dog has not been abroad but visited a forest in Central-Transdanubia (Veszprém County, Hungary), a few days before the surgery. The dog had not been treated with any ectoparasiticides.

The aforementioned forest (coordinates: 46.99, 17.91, situated at an altitude of approximately 311 meters above sea level) was visited on 2025.09.06. late afternoon and on 2025.09.07 from morning until afternoon for the purpose of selective louse fly collection. Flying, unfed specimens were collected using an insect net.

Following these, additional collections have been conducted at various sites across Hungary:

In the Mátra Mountains in Northern Hungary on 2025.09.27 (coordinates: 47.90, 20.01, situated at an altitude of approximately 336 meters above sea level).

In the őrség region in Western Hungary on 2025.10.12 (coordinates: 46.85, 16.51, situated at an altitude of approximately 228 meters above sea level).

Two additional louse flies were found by one of the authors after walking a dog in the Bükk Mountains in Northeastern-Hungary, on 2025.10.13 (coordinates: 48.11, 20.61; approximately 319 meters above sea level). These specimens, already wingless but not yet engorged, were removed from the dog, which showed no signs of dermatological abnormalities.

The shown coordinates are approximate, in all cases, parasite specimens were collected from areas outside of legally protected or designated conservation zones.

The louse flies were identified based on standard morphological keys (Andreani et al., 2019; Maa, 1965; Oboňa et al., 2022; Salvetti et al., 2020) and were also compared to the newly described species *L. andaluciensis* (González et al., 2024; Usai et al., 2025). For species identification, the following morphological attributes were used: the size of the specimens, the number and position of acrostical and postalar bristles, the mesosternal spines and the pregenital sclerite of females).

### Molecular analyses

Four flies (including the *L. fortisetosa* that was initially found on the dog) were chosen for molecular identification: two *L. fortisetosa*, and two *L. cervi*. The surface of louse flies was disinfected by sequential washing for 15 s in detergent, tap water and in distilled water. For the DNA extraction, two legs of each *L. fortisetosa*, and one leg of each *L. cervi* specimens were cut off. DNA was extracted with the QIAamp DNA Mini Kit (Qiagen, Hilden, Germany) according to the manufacturer’s instruction, including an overnight digestion in tissue lysis buffer and Proteinase-K at 56 °C. Extraction controls (tissue lysis buffer) were also processed with the hippoboscid samples to monitor cross-contamination.

The cytochrome *c* oxidase subunit I (*cox* 1) encoding gene was chosen as the target for molecular analysis. The PCR was modified from (Folmer et al., 1994) and amplifies an ∼710-bp-long fragment of the gene. The primers HCO2198 (5′-TAA ACT TCA GGG TGA CCA AAA AAT CA-3′) and LCO1490 (5’-GGT CAACAA ATC ATA AAG ATA TTG G-3’) were used in a reaction volume of 25 μl, containing 1 U (stock 5 U/μl)HotStarTaq Plus DNA Polymerase (Qiagen, Hilden, Germany), 2.5 μl 10 × CoralLoad Reaction buffer (including 15 mM MgCl_2_), 0.5 μl PCR nucleotide Mix (Qiagen, Hilden, Germany) (stock 10 mM), 0.5 μl of each primer (stock 50 μM), 15.8 μl ddH_2_O and 5 μl template DNA. For amplification, an initial denaturation step at 95°C for 5 min was followed by 40 cycles of denaturation at 94°C for 40 s, annealing at 48°C for 1 min and extension at 72°C for 1 min. Final extension was performed at 72°C for 10 min.

A non-template reaction mixture served as the negative control in all PCR analyses. Extraction controls and negative controls remained PCR negative in all tests. The PCR products were purified and sequenced by Eurofins Biomi Ltd. (Gödöllő, Hungary). The BioEdit program was used for quality control and trimming of sequences. The analyses of assembled sequences were performed with BLASTN via GenBank (https://blast.ncbi.nlm.nih.gov). The sequences obtained in the current study were deposited in the GenBank database and are available under accession numbers PX467064-PX467067.

## Results

Based on morphological characteristics the single, specimen collected from the dog on 2025.08.26. was identified as *L. fortisetosa*. (Figure 1).

**Figure 1:**
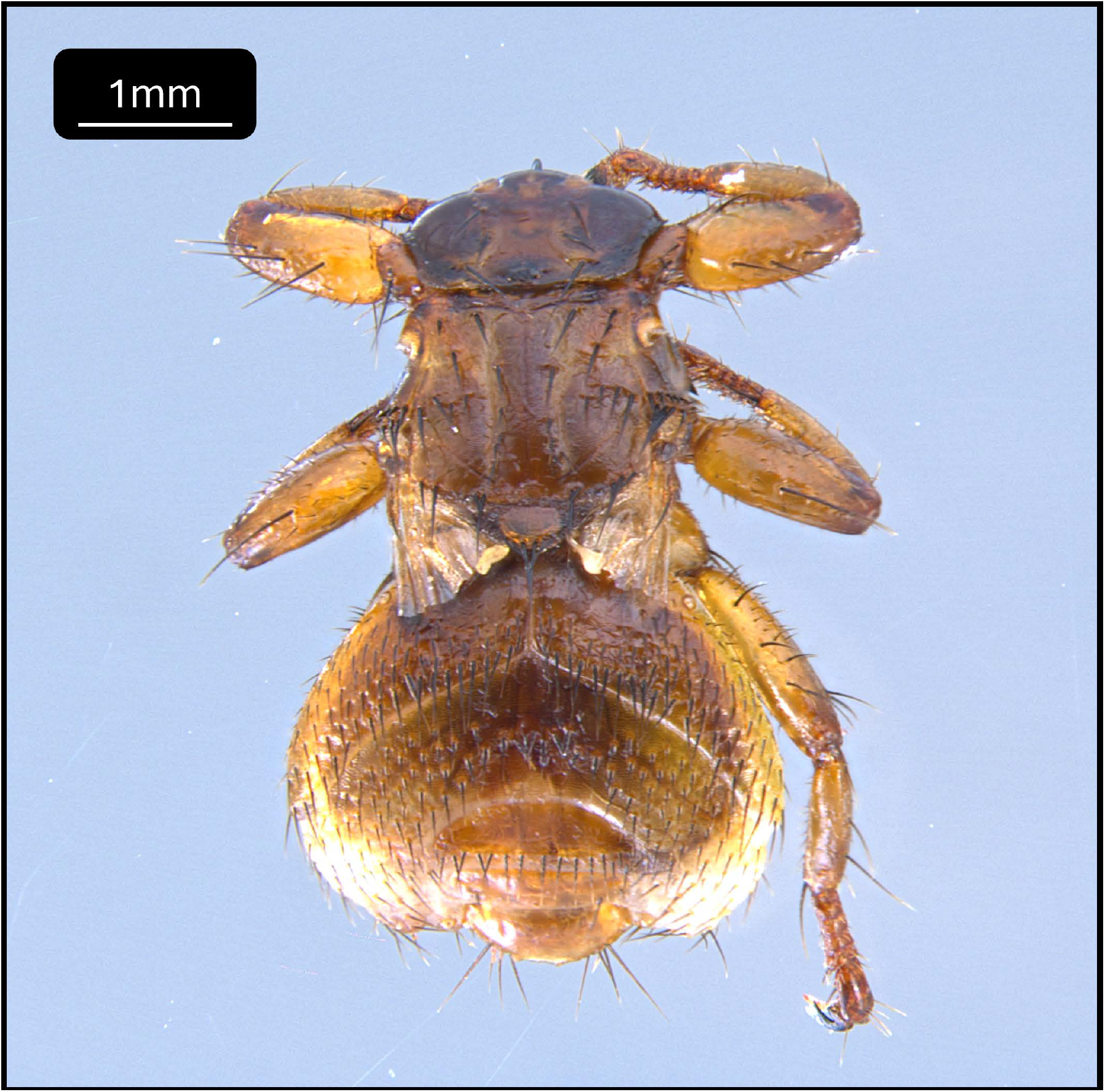
Wingless specimen of *Lipoptena fortisetosa*, removed from a dog.

During the selective louse fly collections on 2025.09.06 and 2025.09.07, 30 specimens of *Lipoptena* spp. were collected. Twenty-three of these were identified as *L. fortisetosa*, while seven as *L. cervi* (Figures 2 and 3).

**Figure 2:**
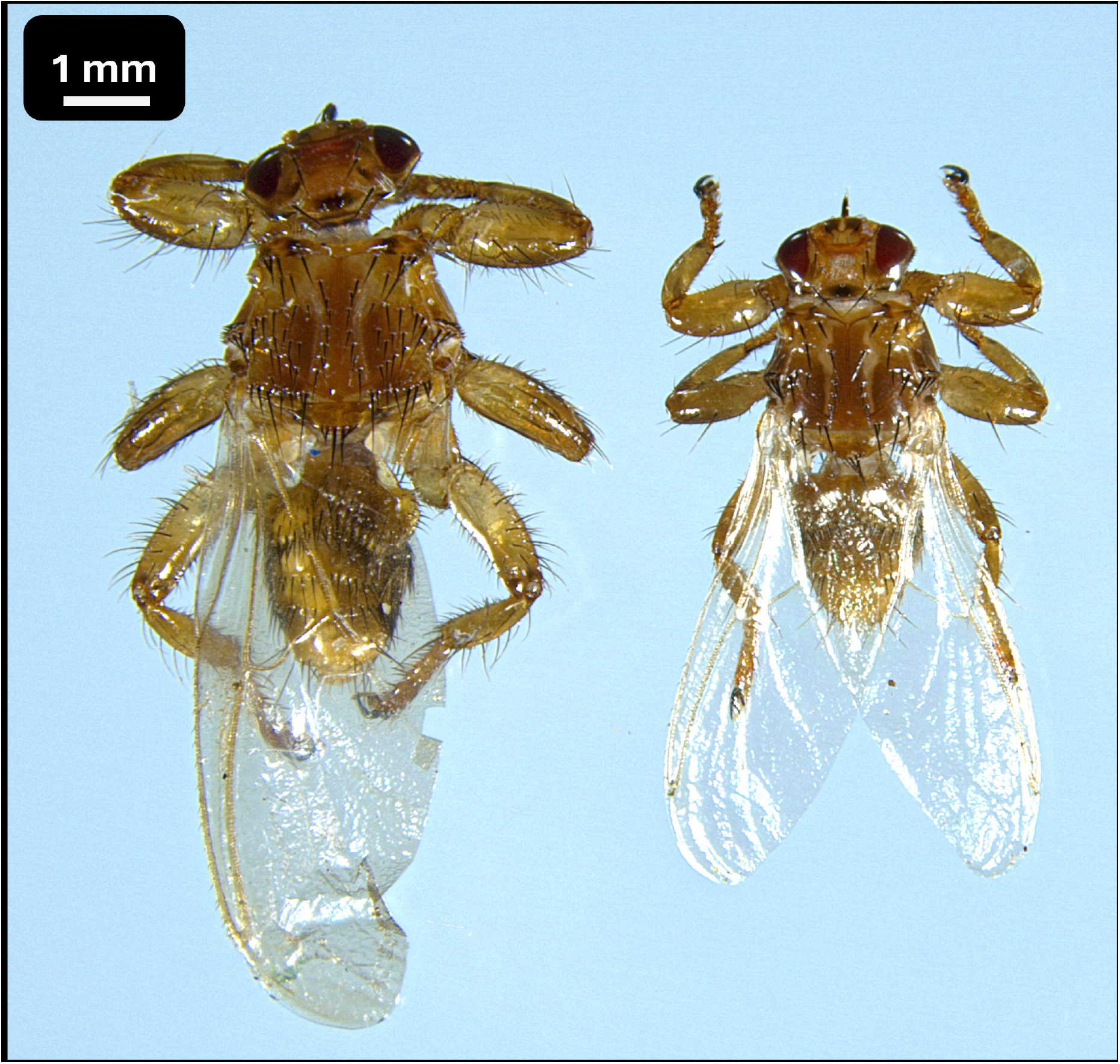
Unfed specimens of *Lipoptena cervi* (right) and *Lipoptena fortisetosa* (left).

**Figure 3:**
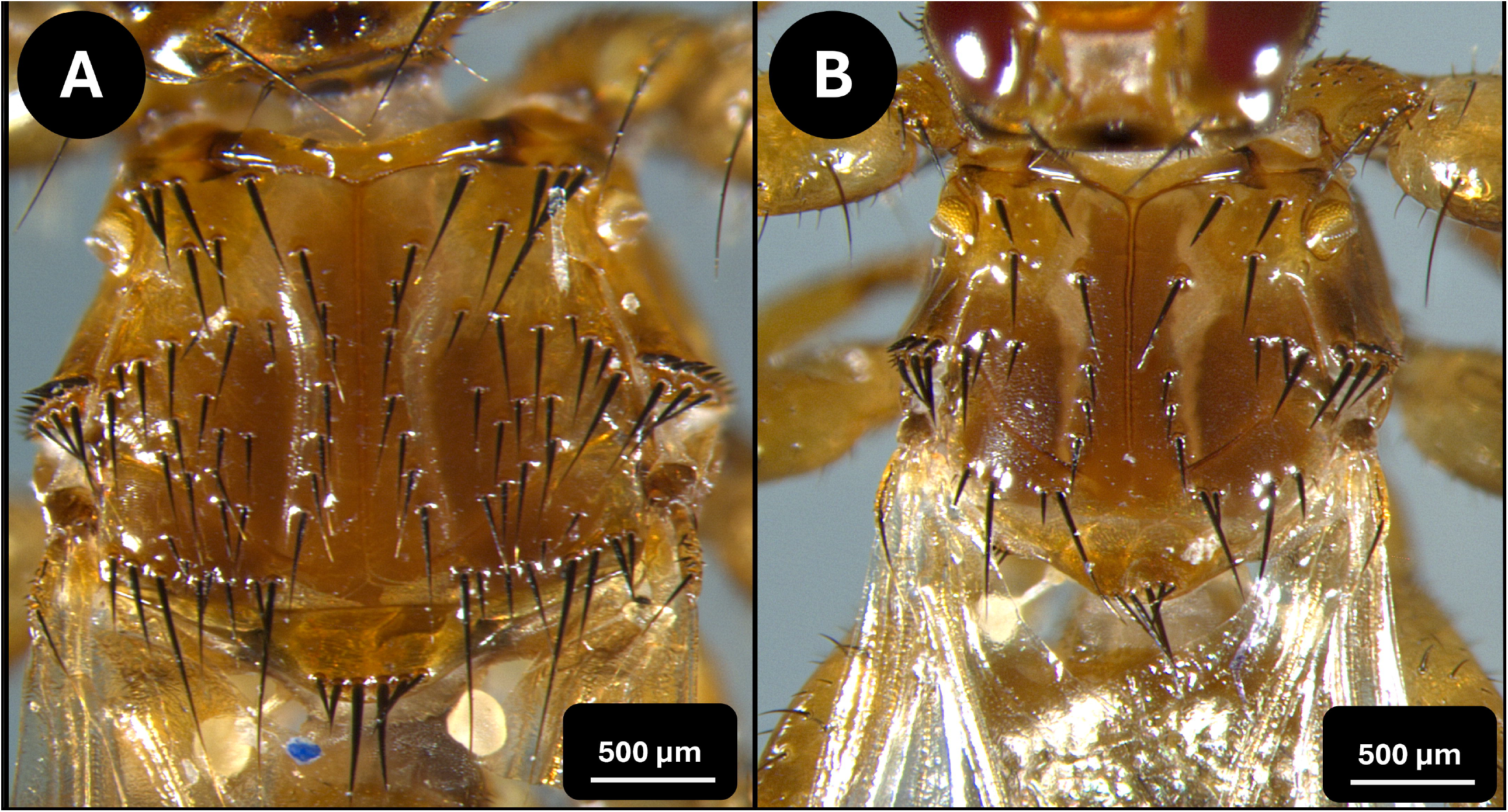
The thorax of *Lipoptena cervi* (A) and the thorax of *Lipoptena fortisetosa* (B).

On 2025. 09.27, five specimens were collected, three of which we identified as *L. fortisetosa*, and two as *L. cervi*.

On 2025.10.12, 45 louse flies were collected, all of them bein*g L. cervi*.

Both flies that were removed from the dog on 2025.10.13 were identified as *L. fortisetosa*.

The results of our molecular analyses based on the *cox*1 gene confirmed our morphological identifications. The flies we identified as *L. fortisetosa* (PX467064 and PX467065) showed 99.85% (652/653 bp) identity to a *L. fortisetosa* specimen from Romania (MK405669) and 100% (643/643 bp) identity to a specimen from Estonia (MN807844). The flies we identified *L. cervi* (PX467066 and PX467067) showed 100% (650/650 bp) and 99.85% identity (649/650 bp) respectively to several *L. cervi* specimens from Scandinavia (KR362270, KR362275)

## Discussion

The occasional presence of *L. fortisetosa* on dogs has been reported in Europe (Kurina et al., 2019; Mihalca et al., 2019; Sokół and Gałęcki, 2017). In addition to this species, *L. cervi* (Hermosilla et al., 2006), *Hippobosca equina* (Maślanko et al., 2022) and *Hippobosca longipennis* (Mihalca et al., 2019) are among the most commonly recorded hippoboscids on dogs. Our report points out, that in contrast to our previous knowledge, dogs are exposed to *L. fortisetosa* in the Hungarian forests, and that the parasite can be passively transported by dogs over considerable distances, contributing to the dissemination of the species. This is supported by the fact that we found this fly on two different dogs. In the initial case, the infested dog had been transported for at least 120 kilometers, and the fly remained on the dog for more than a day, reaching an engorged state. Wing detachment on dogs was also observed on three occasions. This result nuances the previous hypothesis that the species reached Europe through the transportation of deers (Yatsuk et al., 2024), and suggest that pet dogs may play a secondary role in the local dissemination of these ectoparasites. Neither of the dogs exhibited any relevant clinical signs after the removal of the parasites.

Despite the latter facts, it is difficult to determine whether these specimens arrived in Hungary as a result of natural host (deer) migration or through human activity, as, according to new results from Italy, European (including Central European) samples do not differ much from Asian specimens, neither morphologically nor in their molecular characteristics (Andreani et al., 2020). *Lipoptena fortisetosa* was first described in Europe a*s L. parvula* in 1967, in what was then Czechoslovakia (Theodor, 1967). This suggests that the previous absence of *L. fortisetosa* from the Hungarian fauna (Egri and Rigó, 2014; Kaufman, 1988; Papp, 2001) could hardly be interpreted as the true absence of the species in the country, since this absence may be explained (for example) by old, but localized and yet undiscovered populations. Obviously, finding an answer to the question: “How and when *L. fortisetosa* arrived in Europe?” is almost impossible; however, the question itself highlights the importance of the parasite monitoring of imported and transported animals.

To the best of our knowledge, this is the first report confirming the presence and establishment of *L. fortisetosa* in Hungary based on both molecular and morphological evidence. Our data indicate that the species is now widespread in the country. In contrast to previous knowledge, dogs that were walked in forests in central, eastern, and northern Hungary were found to be exposed to this parasite. Furthermore, our findings suggest that *L. fortisetosa* may be passively transported not only by deer but also by dogs.

## Contributions

AGT: Conceptualization, visualization, writing-original draft, writing-review and editing, investigation, validation. AB: writing-original draft, writing-review and editing, investigation. SH, ZsW, BSz, VG, PV: writing-review and editing, investigation. GK: Conceptualization, visualization, writing-original draft, writing-review and editing, investigation, data curation, methodology, validation.

## Acknowledgements

We would like to express our gratitude towards the owner of the infested dog, for her invaluable help.

